# Live cell imaging reveals paclitaxel-induced lysosome motility and function disruption in DRG neurons

**DOI:** 10.64898/2026.05.19.726221

**Authors:** Kathleen Cate Domalogdog, Ishwarya Sankaranarayan, Úrzula Franco-Enzástiga, Juliet M Mwirigi, Shani Mai Nguyen, Diana Tavares-Ferreira, Theodore J Price

## Abstract

Lysosomal trafficking and homeostasis are biological functions that are pivotal for DRG neurons, given their metabolic demands and extremely long axons. Previous studies indicate that lysosomal signaling is altered in a mouse model of chemotherapy-induced peripheral neuropathy (CIPN) and that blocking mitogen activated protein kinase-associated kinase (MNK1/2) signaling can alleviate pain behaviors in CIPN. Here, we investigated lysosome dynamics and lysosome-associated signaling in a mouse model of CIPN induced by paclitaxel (PTX), a chemotherapeutic agent used for various types of cancer. Using spinning disk super-resolution microscope (SPINSR), we demonstrate that PTX treatment *in vivo* causes reduced lysosome motility observed *in* vitro. PTX likewise drives the accumulation of Sequestosome 1 (SQSTM1), also known as P62, in cultured mouse DRG neurons, indicating lysosomal dysfunction in DRG neurons. The transcription factor EB (TFEB), a master regulator of lysosomal biogenesis, was also upregulated in the nucleus of cultured mouse DRG neurons treated with PTX. In line with this, increased lysosomal-associated membrane protein 1 (LAMP1) expression was observed in PTX-treated mice. Given that our previous work demonstrated PTX treatment increases MNK1/2-eIF4E signaling in DRG neurons, we examined whether MNK1/2 inhibition could rescue lysosomal dysfunction. Treatment with Tomivosertib (eFT508), a potent MNK1/2 inhibitor, restored P62 levels in DRG neurons of PTX-treated mice and reduced TFEB in DRG treated *in vitro*. To establish translation relevance, we further show that PTX elevates phosphorylated eiF4E (p-eIF4E) in human DRG neurons, and concurrent eFT508 administration attenuates this effect. Collectively, these findings indicated that PTX disrupts lysosome trafficking and biogenesis, and that MNK inhibition with eFT508 restores lysosomal signaling and can serve as a neuroprotective strategy for CIPN.

## Introduction

Chemotherapy-induced peripheral neuropathy (CIPN) is a common dose-limiting side effect of chemotherapy that may persist long-term(1). Common symptoms typically include pain, allodynia, hyperalgesia, paresthesia, and loss of proprioception(2-4). Paclitaxel (PTX), a chemotherapeutic agent for various types of cancer, such as ovarian and breast cancer(5), causes peripheral neuropathy (CIPN) in 60-70% of patients, reducing their quality of life(4, 6, 7). In cancer, PTX is a potent microtubule-stabilizing agent that inhibits the proliferation of tumor cells(8) by binding to polymerized tubulins (α and β), thereby interrupting the G_2_ phase of the cell cycle(9). However, PTX does not selectively target cancer cells(10). The dorsal root ganglion (DRG) lacks protection from a blood-nerve barrier, and its surrounding capillaries are highly permeable, rendering it vulnerable to PTX accumulation and toxicity(11-13). This microtubule disruption in DRG sensory neurons facilitates axonal degeneration, leading to peripheral neuropathy(14). Previous studies determined that the microtubule dysfunction caused by PTX is prevalent in axons and disrupts fast axonal transport(15-17). This can lead to the compromised neuronal transport of nutrients, neurotransmitters, and organelles(10).

Existing CIPN studies have reported extensively on mitochondrial dysfunction leading to increased reactive oxygen species (ROS) production, which further promotes axonal degeneration(4, 18, 19). It is well known that lysosomes remove dysfunctional mitochondria and other cell debris, but there is evidence that mitochondria and lysosomes have a bidirectional relationship in which the ROS produced by mitochondria can cause lysosomal membrane oxidation, while ruptured lysosomes release cathepsins that promote permeabilization and apoptosis of mitochondria(20-23). This results in a feed-forward loop that exacerbates neuronal degeneration, such as in Parkinson’s disease (24). However, in the context of PTX-induced peripheral neuropathy, there has been very limited information on how PTX affects lysosomes in the DRG.

Lysosomes are highly dynamic organelles with an acidic lumen (pH 4.5–5.0) containing a high content of Ca^2+^ and active hydrolases(25). They are the primary coordinators of macromolecule synthesis and the degradation of damaged organelles(26). Lysosomes are not only degradative organelles, but also nutrient sensors, stress response coordinators, and signal transduction hubs for critical signaling factors such as mechanistic target of rapamycin complex 1 (mTORC1) and TFEB signaling(27-29). Consequently, lysosome integrity is critical for cell homeostasis. Under certain conditions, lysosomes alter their volume/size(30). For instance, lysosomal diameter increases three to five times its original size during nutrient starvation, indicating fusion of lysosomes(31). Moreover, nutrient starvation was found to cause nuclear translocation and calcineurin-mediated dephosphorylation of TFEB, and TFEB activity is strongly linked to nutrient availability, which is mediated by protein phosphorylation(32). mTORC1 phosphorylates TFEB in the presence of nutrients on S211 and S142 serine residues, and phosphorylation of these serines drives the localization of TFEB mainly to the cytosol, and remains inactive there, while mTORC1 inhibition activates TFEB by promoting its translocation to the nucleus (33, 34).

Strong evidence suggests that an interconnected signaling network between mTORC1 and MNK-eIF4E is persistently active in PTX-induced neuropathy, and regulation of translation through MNK⍰eIF4E signaling contributes to the development and persistence of pain hypersensitivity across various preclinical pain models(35-38). Additionally, lysosomal amino acid efflux has been identified as another probable contributor to CIPN, owing to the presence of numerous mRNAs exhibiting increased translation efficiency of proteins that encompass the interface of mTORC signaling with lysosomes(39). Pharmacological inhibition of eIF4E phosphorylation using Tomivosertib (eFT508), an orally bioavailable and potent MNK1/2 inhibitor, has been shown to attenuate increased excitability of nociceptors, suppress mitochondrial dysfunction, and decrease ROS production in preclinical neuropathic pain models(35, 39).

The current study was designed to better understand the effects of PTX on lysosomal dynamics and signaling in mouse DRG neurons. Our findings demonstrate that PTX induces lysosomal dysfunction that is reflected in lysosomal motility and in lysosome-associated signaling. These effects are reversed by inhibition of MNK1/2 signaling. PTX induces increased MNK1/2 signaling in human DRG neurons, pointing out the translatability of MNK1/2 as a target for treatment. These results support a model where coordinated dysfunction of mitochondria and lysosomes causes PTX-induced neuropathy and suggest that interfering with augmented MNK1/2 signaling can restore normal function following PTX treatment.

## METHODS

### Animals

Experiments were conducted in 8-10 weeks C57BL/6J mice purchased from Jackson Laboratory. Animals were used in experiments one week after their arrival at the University of Texas at Dallas animal facilities. Mice with a mutation in eIF4E (S209A, 4EKI mice) that renders eIF4E non-phosphorylatable were a gift from the Sonenberg laboratory at McGill University and were bred at the University of Texas at Dallas. Genotype was confirmed at weaning using DNA from ear clips. Mice were allocated in groups of four per cage in standard, non-enriched cages, under a non-inverted 12-hour light/dark cycle. Animals had free access to food and water *ad libitum*. All animal procedures were approved by the Institutional Animal Care and Use Committee (IACUC) of the University of Texas at Dallas. Experiments were conducted in accordance with the National Institutes of Health Guide for the Care and Use of Laboratory Animals (Publication No. 85-23, revised 1985). Before and during experimentation, animals were monitored for health in accordance with IACUC guidelines at the University of Texas at Dallas.

### Chemicals for *in vivo* treatment

Paclitaxel ((2α,4α,5β,7β,10β,13α)-4,10-Bis(acetiloxi)-13-{[(2R,3S)-3-(benzoilamino)-2-hidroxi-3-fenilpropanoil]oxi}-1,7-dihidroxi-9-oxo-5,20-epoxitax-11-en-2-il benzoate) was purchased from Sigma Aldrich.

### Injections

PTX was dissolved in 1:1 Kolliphor EL (Sigma-Aldrich)/absolute ethanol, then further dissolved in PBS. PTX was administered as previously reported(39, 40). PTX was administered intraperitoneally (i.p.) every other day (4 mg/kg each) for 1 week, for a cumulative dose of 16 mg/kg.

### Behavior

We conducted behavioral experiments in both male and female age-matched C57BL/6J mice. Animals from each cage were randomly assigned to control or treatment groups. Mice were allowed to acclimate to Plexiglas chambers for 1 h before the von Frey test. Behavioral testers were blinded to treatment in all experiments. Mechanical paw withdrawal threshold was measured using the up-down method using calibrated von Frey filaments. The 50% threshold was calculated as previously described(41). For spontaneous pain assessment, we used mouse grimace scoring(42). Grimace score was performed by blinded testers.

### Western blot

Mice were anesthetized with 2.5% isoflurane and decapitated. Lumbar dorsal root ganglia (DRGs) were rapidly dissected, frozen on dry ice, and stored at -80 ºC until further processing. Tissues were homogenized in ice-cold lysis buffer containing 50 mM Tris (pH 7.4), 150 mM NaCl, 1 mM EDTA (pH 8.0), and 1% Triton X-100, supplemented with phosphatase inhibitor cocktail (Sigma-Aldrich, Cat# P8340, Cat# P5726, Cat# P0044) and protease inhibitor (Thermo Scientific, Cat# 78438). Homogenates were centrifuged at 14000 rpm for 10 min at 4 ºC. Supernatants were collected and stored at -80 ºC. Protein concentration was determined using the BCA assay (Thermo Fisher Scientific, Cat. #23225). 10 µg of protein were denatured, separated by SDS-PAGE in 12% gel, and transferred overnight to PVDF membranes (Millipore Sigma, Cat. #IPFL00010). Membranes were blocked for 2 h in 5% bovine serum albumin (BSA) prepared in 1x TTBS (150 mM NaCl, 200 mM Tris, pH 7.4, 0.1% Tween 20), followed by overnight incubation at 4 ºC with primary antibodies in 5% BSA/ 1xTTBS. Rabbit primary antibodies against SQSTM1/P62 (Cell Signaling Technology, Cat. #5114, 1:1000) and GAPDH (catalog #2118; 1:10,000) were purchased from Cell Signaling Technology. Membranes were washed 3 times with 1x TTBS containing 0.1% Tween-20 and incubated for 2 h at room temperature with goat anti-rabbit IgG secondary antibody (Cat. #111-035-008). After 3 washes, protein bands were visualized using the enhanced chemiluminescence detection system (ECL Plus) and quantified by densitometric analysis with Image Lab software (5.2.1, Bio-Rad) and bands were normalized with housekeeping gene GAPDH.

### Immunohistochemistry

Fresh frozen mouse DRGs were sectioned with a cryostat (Leica CM1950) at 20 µm thickness and mounted onto charged slides. Tissues were fixed in 10% formalin for 10 min, followed by 50%, 70%, and 100% ethanol for 5 min each. The slides were allowed to dry at room temperature until the moisture evaporated. Hydrophobic boundaries were drawn around the sections with Pap-pen (ImmEdge Hydrophobic Barrier Pap-Pen, Cat. #H-4000, Vector Laboratories) until dry. Next, sections were blocked with 10% normal goat serum with 0.3% triton X-100 in 0.1 M PB, pH 7.4, for 1 h at room temperature. Sections were then incubated overnight at 4 ºC in blocking solution containing primary antibodies against LAMP1 (Santa Cruz Biotechnology, Cat# sc-19992). Sections were washed with 0.1 M PB, pH 7.4, and incubated with goat anti-rat secondary antibody (ThermoFisher Scientific, Cat# PA1-84761) for 1 h at room temperature. Sections were washed 3 times with 0.1 M PB, dehydrated, coated with Prolong Gold (Cat. #P36390, Invitrogen), and sealed with glass coverslip. Images were captured on the Olympus FV12000 Confocal Microscope System. Images were analyzed using Olympus CellSens Software. Mean gray intensity was calculated by drawing a region of interest around the neuron and normalizing it to the area.

### Primary cell culture of mouse DRG neurons

#### A. Culture protocol for vehicle- and PTX-treated mice in vivo

Mice treated with either vehicle or PTX were anesthetized with 2.5% isoflurane and euthanized by decapitation. On day 10 after PTX or vehicle injection, cervical, thoracic, and lumbar DRGs were collected in HBSS (ThermoFisher, Cat. #14170161). Tissues were digested with 1 mg/mL collagenase A (Roche, Cat.# Sigma, 10103586001) for 25 min at 37 ºC, followed by a second digestion in a 1:1 mixture of 1 mg/mL collagenase D (Sigma, 1188866001) and papain (Roche) for 20 min at 37 ºC. Tissue was triturated in a 1:1 solution of trypsin inhibitor (Roche) and BSA and passed through a 70 µm cell strainer. Cells were collected by centrifugation to form a pellet and resuspended in DMEM/F12 with GlutaMAX (ThermoFisher Scientific) containing 10% fetal bovine serum (FBS; ThermoFisher Scientific), 5 ng/mL NGF, 1% penicillin and streptomycin, and 3 mg/mL 5-fluorouridine with 7 mg/mL. Cells were plated on poly-D-lysine-coated coverslips in well plates. DRG neurons were maintained at 37°C in a humidified atmosphere containing 5% CO_2._ Media change was performed every other day. Independent cohorts of PTX-treated mice were used for cultures to perform Spinning Disk Confocal Imaging and to assess protein expression by western blot and ICC.

#### B. Culture protocol for in vitro treatment of WT mice

Mice at 8 weeks old were anesthetized with 2.5% isoflurane and euthanized by decapitation. Cervical, thoracic, and lumbar DRGs were immediately harvested, and the roots will be cut off, so only the DRG bulb will be placed in HBSS (ThermoFisher, Cat. #14170161) on ice. Tissues were placed in a pre-warmed enzyme solution containing HBSS (ThermoFisher, Cat. #14170161) and 2mg/mL collagenase IV (Worthington Biochemical, Cat# LS004186) and 0.1□mg/mL DNase□I (Worthington Biochemical, Cat#□LS002139) and incubated for a total of 1.5 hrs at 37°C in a gently shaking warm bath and triturated every 30 mins using sterile glass Pasteur pipettes. Once dissociated, cells will be passed through a 70 µm cell strainer (Celltreat, Cat# 229483) and then gently layered onto 1□mL of 15% BSA to form a gradient and centrifuged at 900□×□g for 5□min (acceleration profile□9, deceleration profile□5) at room temperature. The supernatant is then carefully removed, and the dissociated neurons are resuspended in DMEM/F12 with GlutaMAX (ThermoFisher Scientific) containing 1□% penicillin/streptomycin, 1□% N⍰2 Supplement (Thermo Fisher Scientific, Cat#□17502048), 2□% HyClone Fetal Bovine Serum (Thermo Fisher Scientific, Cat# SH3008803IR), 0.15□mg/mL 5⍰fluoro⍰2′⍰deoxyuridine (FRDU) (Sigma⍰Aldrich, Cat#□F0503), and 0.35□mg/mL uridine (Sigma⍰Aldrich, Cat#□U3003). Cells were plated on poly-D-lysine-coated chamber slides. DRG neurons were maintained at 37°C in a humidified atmosphere containing 5% CO_2._ Media change was performed every other day.

### Primary cell culture of human DRG neurons

All human tissue procurement procedures and ethical regulations were approved by the Institutional Review Board at the University of Texas at Dallas under protocol Legacy-MR-15-237. Dorsal root ganglia were procured from organ donors through a collaboration with the Southwest Transplant Alliance, an organ procurement organization (OPO) in Texas. The Southwest Transplant Alliance obtains informed consent for research tissue donation from first-person consent (driver’s license or legally binding document) or from the donor’s legal next of kin. Policies for donor screening and consent are those established by the United Network for Organ Sharing (UNOS). OPOs follow the standards and procedures established by the US Centers for Disease Control (CDC) and are inspected biannually by the Department of Health and Human Services (DHHS). The distribution of donor medical information is in compliance with HIPAA regulations to protect donor privacy.

Lumbar and thoracic human dorsal root ganglia (hDRGs) were recovered from organ donors at Southwest Transplant Alliance (STA) and other local hospitals. hDRGs were transported in aCSF bubbled with CO_2_ and kept on ice. hDRG tissues were transferred to a sterile petri dish containing aCSF and cleaned thoroughly by removing connective tissue, fat, peripheral nerves, and the dural coats to isolate the ganglia. hDRGs were then minced into approximately 1mm sections with sterile scissors and immediately placed in 5□mL of pre⍰warmed (37□°C) enzyme solution containing 1□mg□/mL STEMxyme□1 solution (Worthington Biochemical, Cat# LS004106), 0.1□mg/mL DNase□I (Worthington Biochemical, Cat#□LS002139), and 10□ng/mL recombinant human β⍰NGF (R&D Systems, Cat#□256⍰GF). and tissue was incubated in a conical tube containing the enzyme solution at 37□°C in a gently shaking water bath overnight, but for no more than 8 hrs. Next, tissue was triturated using fire-polished sterile glass Pasteur pipettes with progressively smaller tip diameters until complete digestion was achieved. The resulting cell suspension was filtered through a 100□µm cell strainer (VWR, Cat#□21008-950) and then gently layered onto 3□mL of 10□% BSA to form a gradient and centrifuged at 900□×□g for 5□min (acceleration profile□9, deceleration profile□5) at room temperature. The supernatant were then carefully removed and the dissociated neurons were resuspended in BrainPhys medium (STEMCELL Technologies, Cat#□05790) containing 1□% penicillin/streptomycin, 1□% GlutaMAX (United States Biological, Cat#□235242), 2□% NeuroCult SM1 (STEMCELL Technologies, Cat#□05711), 1□% N⍰2 Supplement (Thermo Fisher Scientific, Cat#□17502048), 2□% HyClone Fetal Bovine Serum (Thermo Fisher Scientific, Cat# SH3008803IR), 25□ng/mL recombinant human β⍰NGF (R&D Systems, Cat#□256⍰GF), 0.15□mg/mL 5⍰fluoro⍰2′⍰deoxyuridine (FRDU) (Sigma⍰Aldrich, Cat#□F0503), and 0.35□mg/mL uridine (Sigma⍰Aldrich, Cat#LU3003). hDRG neurons were then seeded in Poly-D-lysine (PDL, Sigma Aldrich, Cat# P7405) coated well-plate and incubated for 3 hours at 37°C. Once neurons were adhered, additional culture media were added, and the culture was maintained at 37°C with 5% CO_2_. The media was changed every other day.

### Immunocytochemistry

#### A. ICC protocol for DRG neurons from vehicle- and PTX-treated mice in vivo

On day *in vitro* (DIV) 4, DRG neurons from vehicle- or PTX-treated groups were incubated with 75nM LysoTracker Red DND-99 (Invitrogen, Cat# L7528) for 1 hour, washed with 1XPBS, fixed in 10% formalin at room temperature for 10 minutes, and gently rinsed with 1XPBS. The cells were then blocked with 10% NGS and 0.3% Triton X-100 in 0.1 PB for 1 hour. The cells were then incubated overnight with anti-Lamp1 antibody (Santa Cruz Biotechnology, Cat# sc-19992), washed three times, and incubated with goat anti-rat secondary antibody (ThermoFisher Scientific, Cat# PA1-84761) for 1 hr at room temperature. Images were analyzed using Olympus CellSens Software. Mean gray intensities were calculated by drawing a region of interest around the neuron and normalizing it to the area of the vehicle.

#### B. ICC protocol for DRG neurons from WT mice treated in vitro

On DIV7, mDRG cultures were treated with PTX (Selleck Chemicals, Cat# S1150) or PTX + eFT508 (MedChem Express, Cat#HY-100022) for 24 hrs. Both compounds were reconstituted in DMSO to 10 mM and stored at -80°C for long-term storage. After the 24h treatment, cells were fixed with ice-cold 10% formalin for 10 minutes and washed with 1XPBS. The cells were then blocked with 10% NGS and 0.3% Triton X-100 in 1XPBS for 1 hour. Followed by overnight incubation with rabbit primary antibody against TFEB (Proteintech, Cat# 13372-1-AP) and chicken primary antibody against peripherin (Encor biotechnology, Cat# CPCA-Peri), washed three times, and incubated with DAPI, goat anti-rabbit (Invitrogen, Cat# A21245) and anti-chicken (Invitrogen, Cat# A-11039) secondary antibodies for 1 hr at room temperature. Images were analyzed using Olympus CellSens Software. Mean gray intensities of nuclei were acquired by drawing a region of interest around the nuclei of neurons using DAPI as a reference, and the mean gray intensity of cytosol was determined by calculating the difference of the integrated intensities of the whole neuronal body and the nucleus divided by the difference of the whole neuronal body area and the nucleus area. All mean gray intensity values are then normalized to the vehicle.

#### C. ICC protocol for human DRG neurons treated in vitro

On DIV7, hDRG cultures were treated with PTX (Selleck Chemicals, Cat# S1150) or PTX+ eFT508 (MedChem Express, Cat#HY-100022) for 24 hrs. Both compounds were reconstituted in DMSO to 10 mM and stored at -80°C for long-term storage. After the 24h treatment, cells were fixed with ice-cold 10% formalin for 10 minutes and washed with 1XPBS. The cells were then blocked with 10% NGS and 0.3% Triton X-100 in 1XPBS for 1 hour. Followed by overnight incubation with rabbit primary antibody against p-eIF4E^S209^ (Abcam, Cat #ab76256) and chicken primary antibody against peripherin (Encor biotechnology, Cat# CPCA-Peri), washed three times, and incubated with DAPI, goat anti-rabbit (Invitrogen, Cat# A21245) and anti-chicken (Invitrogen, Cat# A-11039) secondary antibodies for 1 hr at room temperature. Images were analyzed using Olympus CellSens Software. Mean gray intensities were calculated by drawing a region of interest around the neuron and normalizing it to the vehicle.

### Live Cell Imaging

Mouse DRG neurons from vehicle- and PTX-treated mice on DIV4 *in vitro* were stained with 75nM LysoTracker Red DND-99 (Invitrogen, Cat# L7528) for one hour. The dye-containing media were replaced with fresh media before imaging. The cells were stabilized for at least 20 minutes in a stage-top incubator chamber at 37°C with 5% CO_2_. Live-cell imaging was performed using a SPINSR spinning disk super-resolution microscope, and Z-stack images were acquired every 5 seconds for 5 minutes at 100X magnification. ImageJ software was utilized for analysis.

### Data Analysis

Data were analyzed with Graphpad Prism V9 (Graphpad, San Diego, CA). Data are shown as mean ± standard error of the mean (SEM). Violin plots were used in selected analyses to visualize data distribution and individual measurements. For experiments comparing vehicle vs. PTX effects, a student’s *t*-test was used. One-way ANOVA was used to analyze the effects of vehicle, PTX, and PTX plus eFT508, while two-way ANOVA was used for experiments with a treatment by time or similar design. *Post hoc* tests following ANOVA analyses are described in the figure legends.

## Results

CIPN, a debilitating side effect of agents such as PTX, currently lacks an effective remedy for prevention or treatment(1). Moreover, the underlying mechanisms of CIPN remain underexplored. Given previous reports of lysosomal dysfunction caused by PTX in DRG neurons(39), we hypothesized that lysosomal trafficking in DRG neurons may be disrupted by PTX treatment. We injected PTX or vehicle into C57BL/6 mice (**Fig. 1A**) as previousl described(39, 40). Using this administration scheme, we observed tactile hypersensitivity and heightened grimacing in male and female mice (**Sup. Fig. 1**). We used a SPINSR spinning disk super-resolution microscope to assess lysosome motility in the cell body of cultured DRG neurons from PTX-treated mice (**Fig. 1B, Video**). We found that the average speed of lysosomes in neurons from PTX⍰treated mice was reduced compared to neurons from vehicle⍰treated mice (**Fig. 1C**). These results suggest that PTX exposure impairs lysosomal motility, potentially contributing to CIPN.

**Figure 1.**
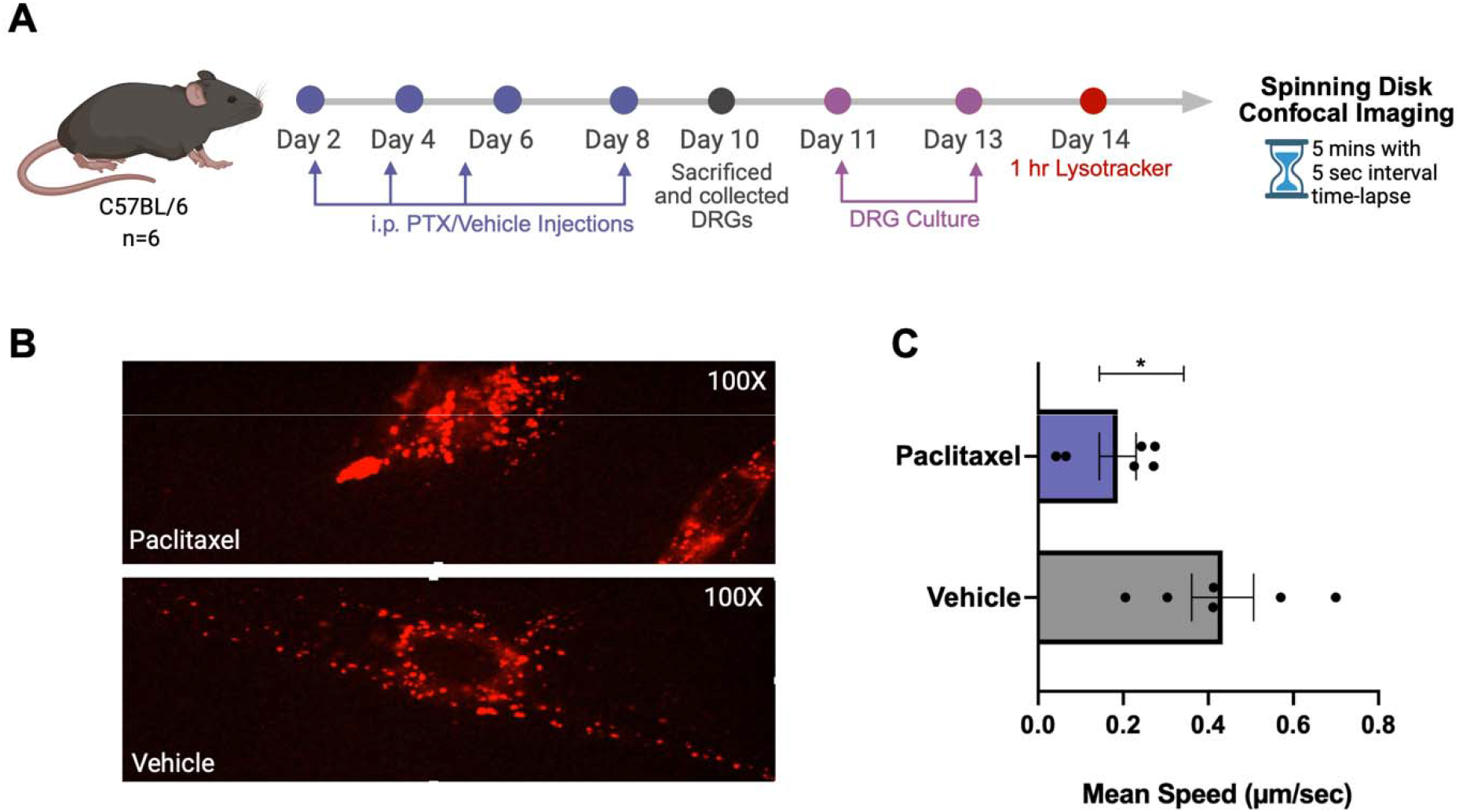
PTX treatment disrupts the motility of lysosomes and drives accumulation to the cytosol of mouse DRG neurons. **A**. Diagram illustrating the experimental timeline for lysosome tracking. C57BL/6 mice received either vehicle or 4 mg/kg PTX via intraperitoneal injections every other day for a total dose of 16 mg/kg, followed by DRG dissection and culture. Live-cell imaging of lysosomes was conducted on day 14 using the SPINSR spinning-disk super-resolution microscope. **B**. Representative time-lapse video of mouse DRG neurons stained with Lysotracker shows reduced motility and accumulation of lysosomes in the soma. **C**. PTX injection significantly reduces the motility of lysosomes in mouse DRGs neurons compared to vehicle. Data are presented as mean ± SEM. Statistical significance was determined using an unpaired two-tailed Student’s t-test. *p < 0.05

Lysosomes are not static organelles; they dynamically adapt to extracellular and intracellular stimuli that alter their size, position, and quantity(43). Increased lysosomal size also negatively affects exocytosis(44), and there is an inverse correlation between lysosomal size and diffusive movement(45). Hence, our findings prompted us to examine whether PTX injection would also trigger these changes. To assess this, we determined the average area of lysosomes in animals treated with PTX or vehicle (**Fig. 2A-B**). We found that the average area of the lysosomes increased in neurons from PTX -treated animals (**Fig. 2C**), suggesting that this chemotherapeutic agent enlarges the lysosomal compartment, potentially leading to lysosomal stress or even lysophagy. We used both Lysotracker Red DND-99 and LAMP1 to stain lysosomes, since co-labeling is a reliable strategy for distinguishing lysosomes from other acidic compartments, such as late endosomes (**Fig. 2B**). The zoomed images of the cell body of the DRG neurons from PTX-mice show swelling and the colocalization of LAMP1 antibody and Lysotracker dye, as pointed by the white arrows, suggesting that enlarged lysosomes are found in PTX-treated mice.

**Figure 2.**
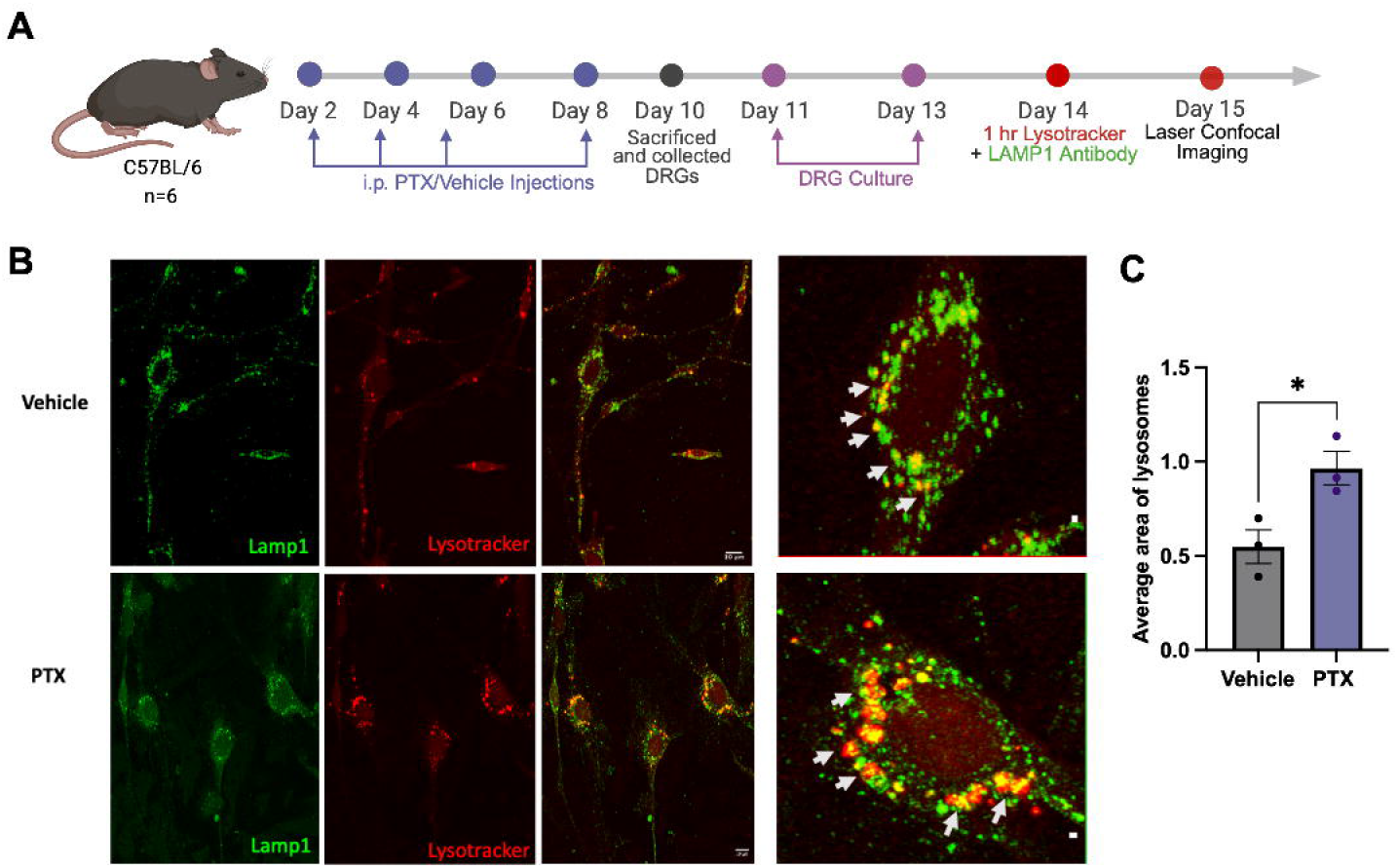
Lysosomes of DRG neurons from PTX-treated mice significantly increased in size. **A**. Experimental timeline showing the frequency of parallel intraperitoneal injections of either 4 mg/kg PTX or vehicle administered to C57BL/6 mice. DRGs were harvested from the treated mice, cultured for 3 days, and stained with both Lysotracker dye and an LAMP1 antibody. Laser confocal imaging was performed on day 15 after conducting immunocytochemistry. **B**. Representative 40X images of mouse DRG neurons showing lysosomes stained with both LAMP1 antibody (green) and Lysotracker dye (red) in vehicle and PTX treatment conditions. Contrast uniformly adjusted for presentation purposes only. White arrows pointing to lysosomes co-labeled with LAMP1 and Lysotracker. **C**. Lysosomes in the DRG neurons of PTX-treated mice show a significant increase in their area compared to vehicle-treated mice, as determined by the area of lysosomes co-labeled with LAMP1 and Lysotracker, as indicated by the arrows. Data are presented as mean ± SEM. Image scale bar = 20 µm. Comparison between groups was analyzed using an unpaired two-tailed Student’s t-test. *p < 0.05

Since we observed a decrease in lysosomal speed as well as their enlargement, we decided to examine lysosomal integrity in animals injected with PTX. The loss of lysosomal integrity can activate cell death pathways. Therefore, strict mechanisms of lysosomal quality control are activated to protect cells. Emerging models of lysosomal quality control indicate that upon lysosomal damage, p62/SQSTM1 is recruited to ubiquitinated cargo on damaged lysosomes to facilitate their autophagic clearance (lysophagy); importantly, because p62 is itself degraded through this process, its accumulation serves as a reliable indicator of impaired autophagic flux and lysosomal degradative capacity(46) (47). We hypothesized that upon PTX -induced lysosomal stress, p62 would be activated. We found that PTX injection leads to p62 accumulation in DRG **(Fig. 3)**. p62 transcription is modulated by autophagy and lysosomal regulator transcription factor EB (TFEB)(48, 49), whose nuclear translocation depends on mTORC1(50). This mTORC1 detachment from the lysosome during nutrient-starved states strongly triggers the initiation of autophagy(51). Our group has demonstrated that an underlying mechanism of PTX-induced CIPN in mouse DRG neurons is enhanced mTORC1 signaling at the lysosomal surface that can be attenuated by blockade of MNK1-eIF4E signaling(39). Therefore, we tested whether a MNK1/2 inhibition would modulate the p62 pathway. We observed that the combination of PTX with eFT508 reverses the p62 increase in DRG neurons cultured from mice treated with PTX *in vivo* (**Fig. 3A-B**). eFT508 alone decreased p62 levels in DRG neurons.

**Figure 3.**
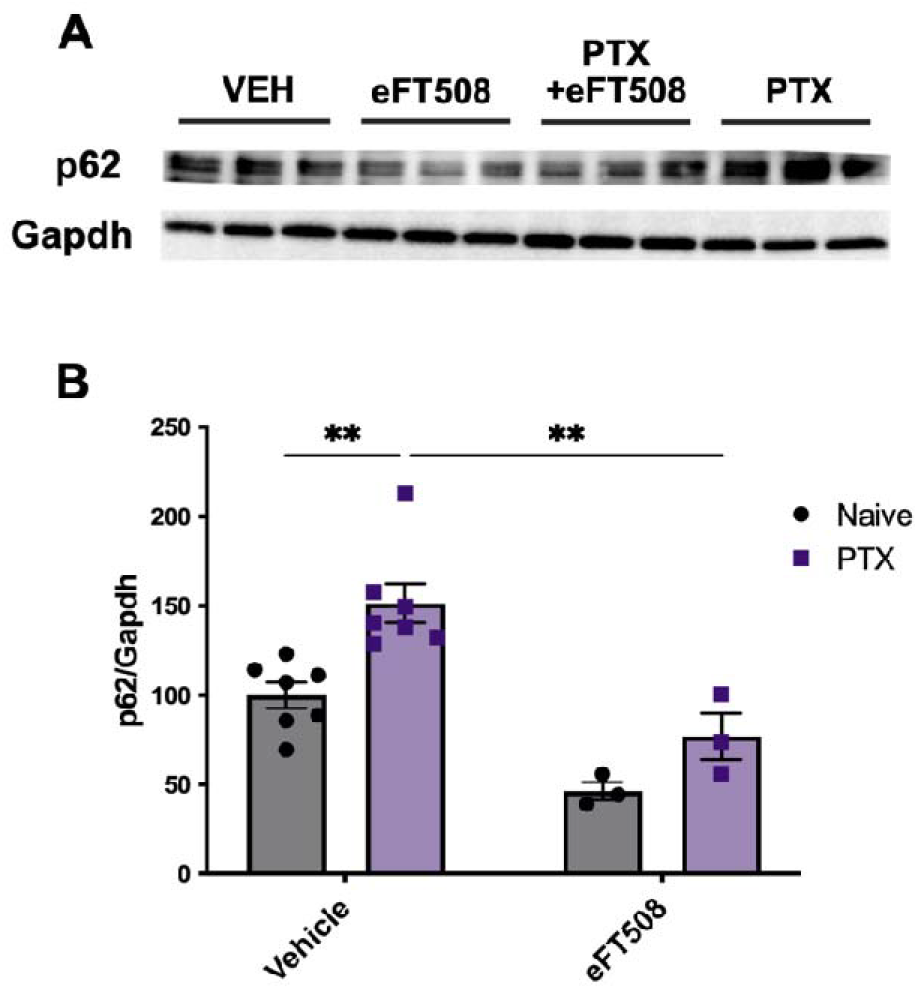
PTX treatment increases p62 in mouse DRGs, while eFT508 reverses this effect. **A**. Representative western images showing p62 bands in blot DRGs of PTX-injected mice or vehicle, plus eFT508. **B**. Western blot analy is of p62 in DRGs of WT mice (n = 3-7 per group) treated with PTX and vehicle, plus eFT508, or eFT508 only. Data are presented as mean ± SEM \*\**p* < 0.01 as determined by two-way ANOVA followed by Bonferroni’s multiple comparisons test.

TFEB, the master transcriptional regulator of lysosomal biogenesis and stress response, has been proposed as the driver of the regeneration of the lysosomal pool(52). Its activation has emerged as an important player in lysosomal damage and repair mechanisms(33, 34, 50). P62 modulation in PTX-treated animals prompted us to assess TFEB expression in mouse DRG cultures treated with PTX *in vitro* (**Fig. 4A**). We found that TFEB is activated by PTX in these cells, specifically leading to increased expression in the nucleus (**Fig. 4A-B**). Moreover, when administered together with eFT508, we observed a significant decrease in nuclear TFEB expression, along with a notable reduction in the cytosol (**Fig. 4C**).

**Figure 4.**
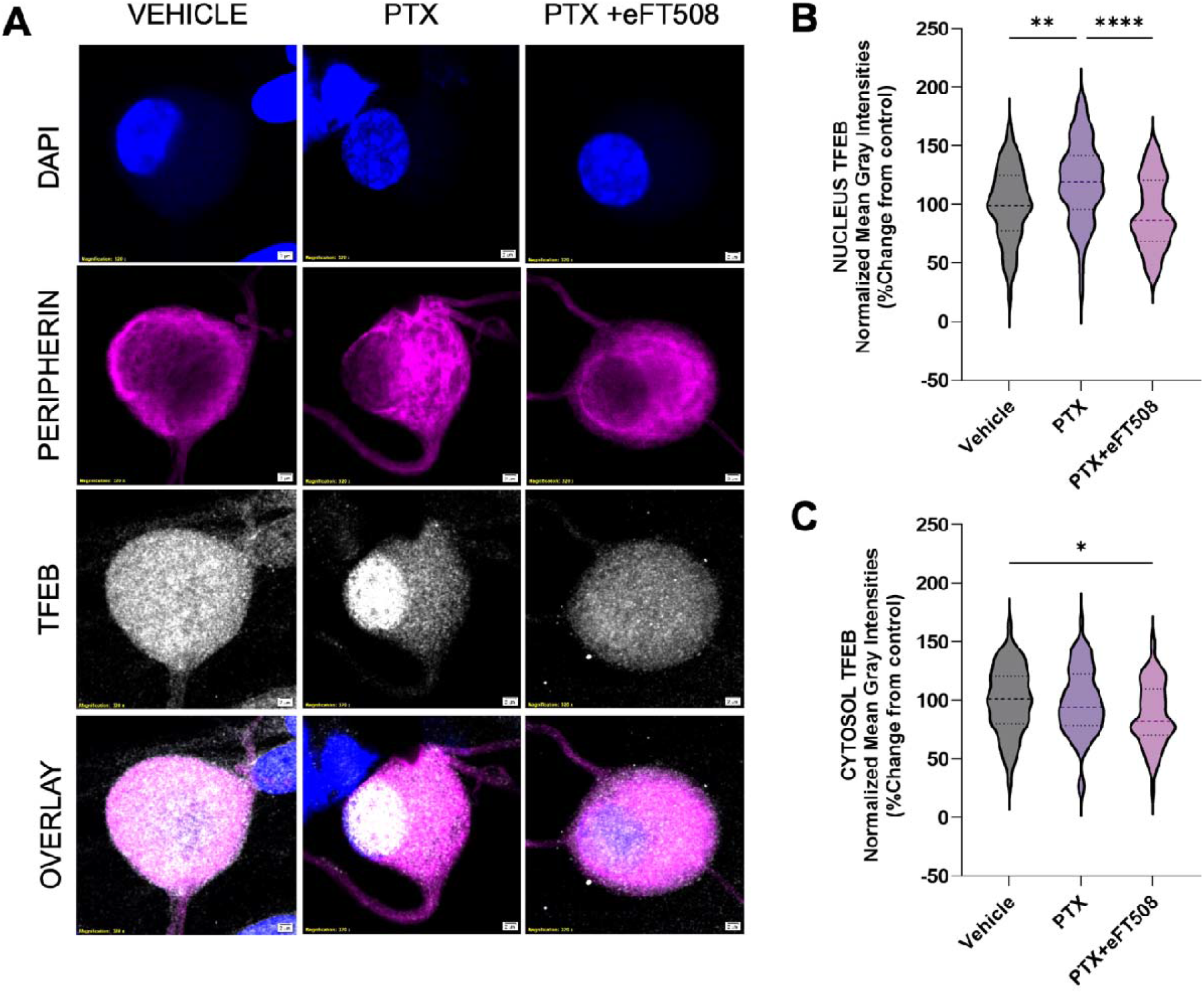
*In vitro* PTX treatment increases TFEB and translocates it to the nucleus of mouse DRG neurons, and concurrent administration of eFT508 exhibits the reversal of this effect. **A**. Representative confocal images of mouse DRG neurons under vehicle (0.002% DMSO), 100 nM PTX, and 100 nM PTX + 100 nM eFT508 conditions showing TFEB expression in gray, peripherin (magenta) to identify neurons, and DAPI (blue) to stain nuclei. Images were captured using a 40x objective lens with an additional 8x digital zoom to highlight the nucleus and cytosol. Contrast has been uniformly adjusted for presentation purposes only. **B**. Normalized mean gray intensities reveals a significant increase in nuclear TFEB expression after 24 hours of PTX treatment, which was attenuated by concurrent administration of eFT508. Interestingly, there is also a marked decrease in cytosolic TFEB expression, as shown in **C**. Mouse DRG cultures were prepared independently from two mice. We analyzed 58, 60, and 74 neurons for vehicle, 100 nM PTX, and 100 nM PTX + eFT508 conditions, respectively. Data are presented as violin plots showing the distribution of individual measurements. Image scale bar = 2 μm. One-way ANOVA with Tukey’s multiple comparisons test *p < 0.05, \*\**p* < 0.01, \*\*\*\**p* < 0.0001.

The experiment described above suggests that PTX treatment increases TFEB signaling in the nucleus, an effect which would be expected to increase the number of lysosomes and autophagosomes(51). TFEB nuclear signaling governs the coordinated lysosomal expression and regulation (CLEAR) network and activates its downstream target genes, such as LAMP1(48, 53, 54). LAMP1 is also an important marker of lysosome formation. Therefore, we hypothesized that PTX treatment would increase LAMP1 expression in the DRG, and that this effect might be attenuated by disrupting MNK1/2 signaling. To test this, we treated mice with PTX using the same treatment regimen described above. We did this experiment in WT and 4EKI mice. We chose to genetically reduce MNK1/2-eIF4E signaling so that we did not have to treat mice with daily eFT508 treatments concomitantly with PTX treatments. Our findings show that PTX treatment increased LAMP1 expression in WT PTX-treated mice, but this effect was completely lost in the 4EKI mice (**Fig. 5A-B**). This supports the conclusion that the MNK1/2 pathway orchestrates lysosomal homeostasis pathways activated by PTX-induced stress, and that there is heightened lysosome production and/or lysosomal retention in the neuronal cell body occurring with PTX administration.

**Figure 5.**
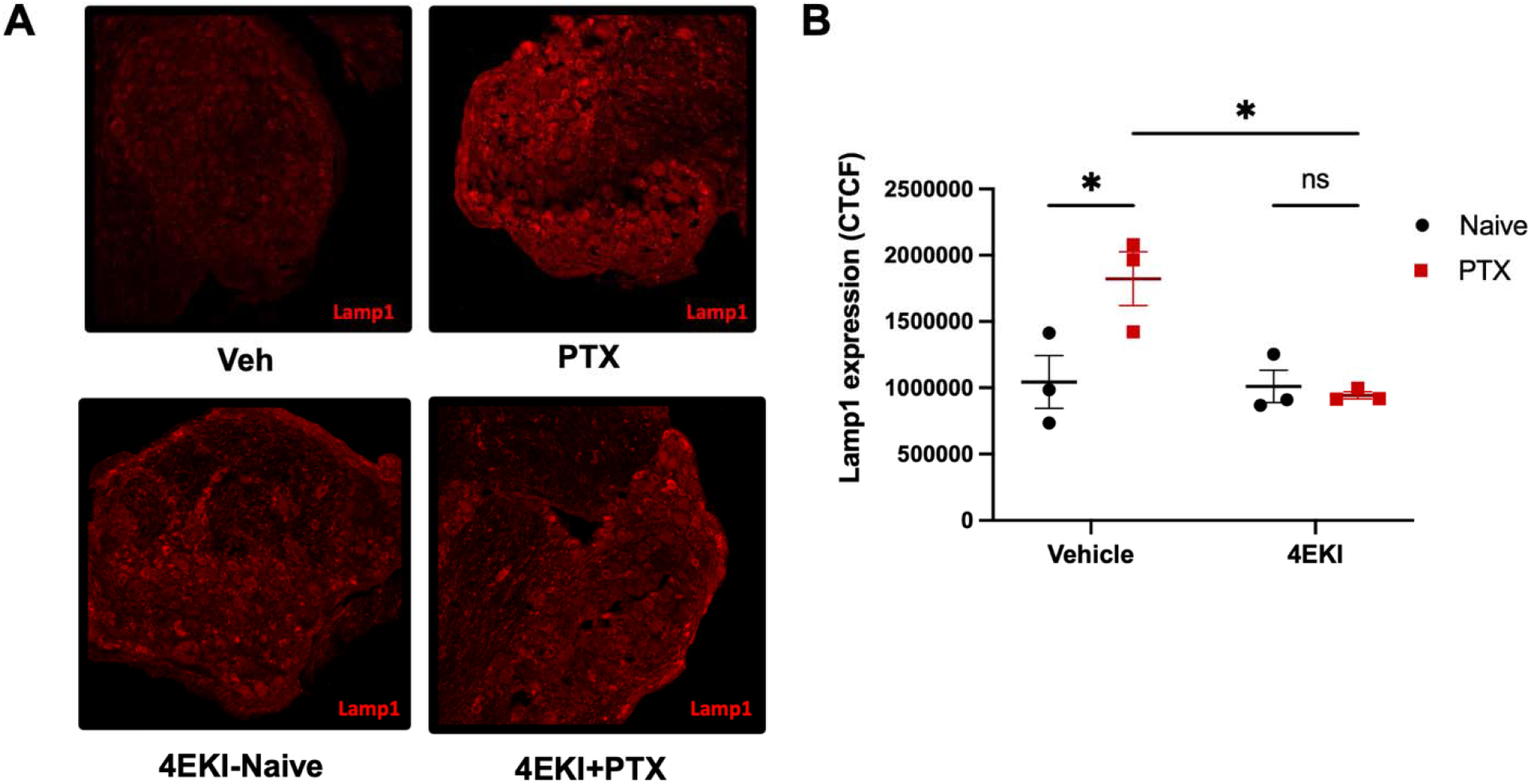
LAMP1 expression is increased by PTX and this effect is lost in 4EKI mice. **A**. Representative images of WT and 4EKI mouse DRG neurons showing LAMP1 expression (red) under vehicle or PTX-treated conditions. **B**. 4EKI mice treated with PTX exhibit significant reversal of LAMP1 expression in the DRG compared to PTX-treated WT mice. Data are presented as mean ± SEM from three independent experiments. Statistical analysis was performed using two-way ANOVA with Sidak’s multiple comparisons test. *p < 0.05, scale bar = 20 μm.

Our present findings, as well as previous studies, support the conclusion that MNK-eIF4E signaling is involved in CIPN development and maintenance in rodent models(35-38). However, this has not been tested in any human model system. To assess whether MNK1/2 signaling is induced in human DRG (hDRG) neurons with PTX treatment we cultured hDRG recovered from organ donors and treated these neurons with PTX and then measured MNK1/2 signaling by examining eIF4E phosphorylation (**Fig 6A**). As shown in Fig. **6B** and **6C**, PTX significantly increased the phosphorylation of eIF4E, and concurrent administration with eFT508 reversed this effect.

**Figure 6.**
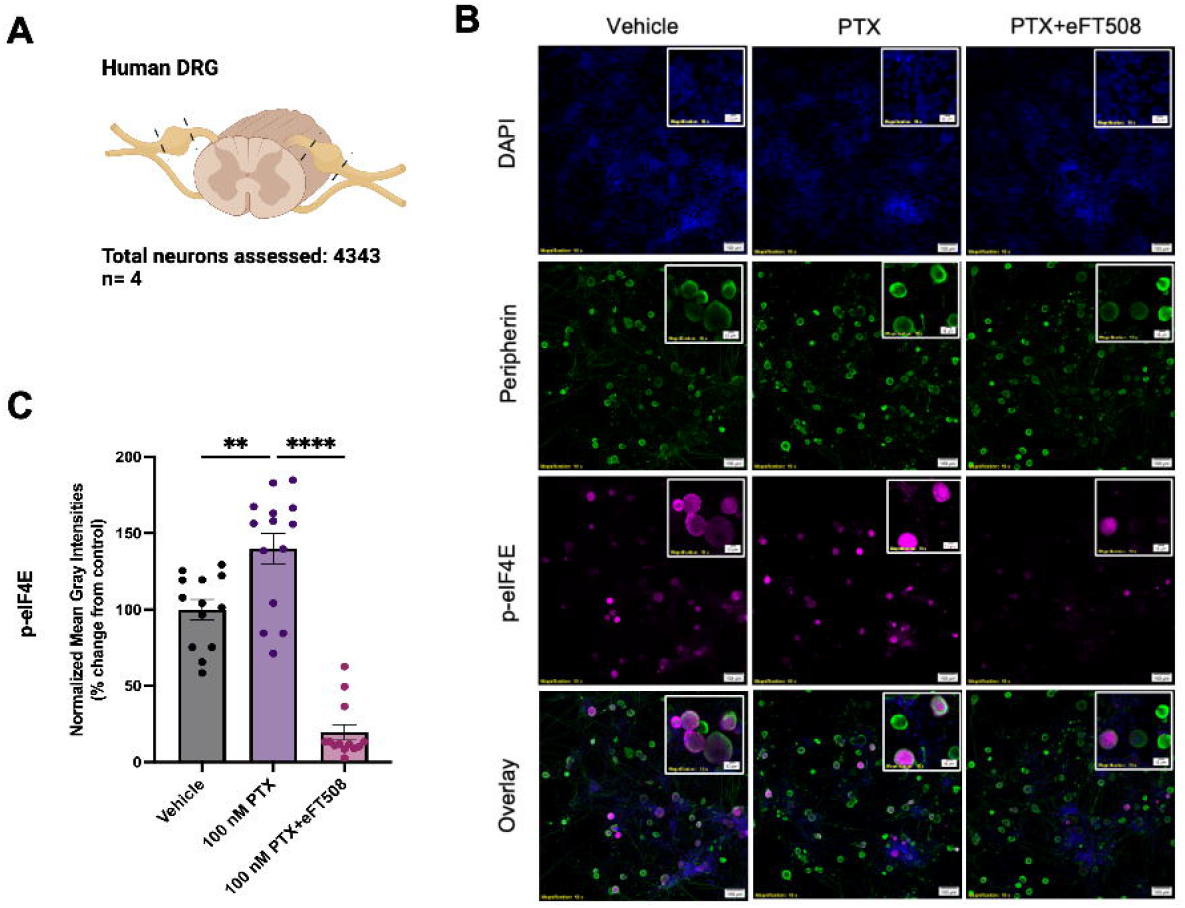
PTX increases the phosphorylation of eIF4E in human DRG neurons. **A**. Schematic diagram of a human spinal cord section with DRGs. Lines indicate where the nerves are cut to isolate the DRG. Details on the number of neurons assessed and the number of biological replicates are included. **B**. Representative 10X images of human DRG neurons under various experimental conditions showing p-eIF4E expression in magenta, peripherin (green) to identify neurons, and DAPI (blue) to stain nuclei. Contrast has been uniformly adjusted for presentation purposes only. **C**. hDRG neurons treated with 100 nM PTX *in vitro* for 24 hours showed significant upregulation of p-eIF4E, and concurrent treatment with 100 nM eFT508 demonstrated robust inhibition of p-eIF4E after 24 hours. DRG cultures were independently prepared from 4 organ donors. N = 13, 14, 13 technical replicates for Vehicle (0.002% DMSO), 100 nM PTX, and 100 nM PTX + eFT508 conditions, respectively. Data are presented as mean ± SEM. Image scale bar = 100 μm. One-way ANOVA with Tukey’s multiple comparisons test \*\**p* < 0.01, \*\*\*\**p* < 0.0001.

## Discussion

Our current findings show that the anti-neoplastic chemotherapeutic agent PTX dysregulates lysosomal homeostasis in DRG neurons by impeding lysosomal motility, altering lysosomal morphology, and regulating transcriptional signaling related to lysosomal biogenesis and function. Connected to this latter point, we demonstrated that PTX drives the accumulation of p62, increases LAMP1 expression, and promotes TFEB translocation to the nucleus, suggesting nutrient deficits in DRG neurons caused by PTX treatment and the activation of autophagy and mTORC1 pathways, consistent with previous findings(39). Moreover, we demonstrated that PTX significantly upregulates eIF4E phosphorylation in human DRG neurons, supporting previous findings in preclinical models of CIPN(35-38). Our study also further supports the finding that the potent MNK1/2 inhibitor, eFT508, acts as a neuroprotective agent, specifically protecting lysosomes from stress in the presence of PTX. This was exemplified by significant decreases in p62, LAMP1, and p-eIF4E expression, as well as in the cytoplasmic localization of TFEB, in DRG neurons concurrently treated with PTX and eFT508.

PTX is a microtubule-stabilizing agent that binds to β-tubulin on the luminal side of microtubules and strengthens longitudinal contacts, which promotes their assembly and stability(55). The consequence of increased microtubule stabilization in neurons is impaired axonal transport(14-17). This was supported by our results showing that lysosomal trafficking speed is slowed by PTX, indicating a deficiency in nutrient delivery and the clearance of damaged organelles and other cellular waste. This motility disruption might also contribute to the swelling of lysosomes that we observed in our experiments. A recent study showed that under stress conditions, lysosomes vacuolate to accommodate the increased contents through the recruitment of PDZ domain-containing protein 8 (PDZD8), a tunnel-like lipid transfer protein acting as a bridge between the endoplasmic reticulum and lysosomes that enables this lysosomal membrane expansion(56).

In this work, we show that PTX increases eIF4E phosphorylation in human DRG neurons and eFT508 attenuates this effect. This posttranslational modification controls the translation of a group of mRNAs involved in neuronal plasticity and inflammation(57), aligning with increasing evidence for the role of MNK1/2-eIF4E in pain across preclinical models(39, 58) and in human DRG neuron experiments(59). Former work from our group showed that the chemotherapeutic agent vinorelbine, from the vinca alkaloid family, also causes pain through engagement of the MNK1/2-eIF4E pathway via activation of cGAS-STING-IFN signaling(38). Interestingly, lysosomal damage can promote the activation of cGAS-STING, rendering this pathway a potential axis for chemotherapy-induced damage that may be common between vinorelbine and PTX (60). Further work will be needed to test this hypothesis, however, our work does support the idea that MNK1/2 signaling is a key regulator of CIPN.

There are several limitations to our work. First, while our findings support the notion that lysosomal dysfunction is an important feature of CIPN, *in vivo* assessment of lysosomal dynamics would provide stronger support for this conclusion. This work would be technically challenging, and our view is that our *in vitro* experiments set the stage for how those experiments could be done. Second, we did not assess TFEB-induced transcriptional changes using RNA-sequencing approaches. Our findings with LAMP1 suggest TFEB-induced transcriptional changes but this should be assessed in the future using single nucleus RNA-seq or similar approaches. Finally, we only assessed a single endpoint in our hDRG work. We plan to conduct studies to assess lysosomal live cell dynamics in these neurons upon chemotherapeutic treatment in the future.

## Supporting information

Supplementary Video Vehicle

Supplementary Video Paclitaxel

## Acknowledgments

This research was supported by the National Institute of Neurological Disorders and Stroke of the National Institutes of Health grant NS065326 and through the PRECISION Human Pain Network (RRID:SCR_025458), part of the NIH HEAL Initiative (https://heal.nih.gov/) under award number U19NS130608 to TJP. The content is solely the responsibility of the authors and does not necessarily represent the official views of the National Institutes of Health.

## Notes

**Conflict of Interest Statement:** TJP is a founder of and holds equity in 4E Therapeutics, PARMedics, NuvoNuro, Nerveli, and Ted and Greg’s. The other authors declare no conflicts of interest.

### Competing Interest Statement

TJP is a founder of and holds equity in 4E Therapeutics, PARMedics, NuvoNuro, Nerveli, and Ted and Gregs. The other authors declare no conflicts of interest.

