## Supplementary Figures for "Live cell imaging reveals paclitaxel-induced lysosome motility and function disruption in DRG neurons"

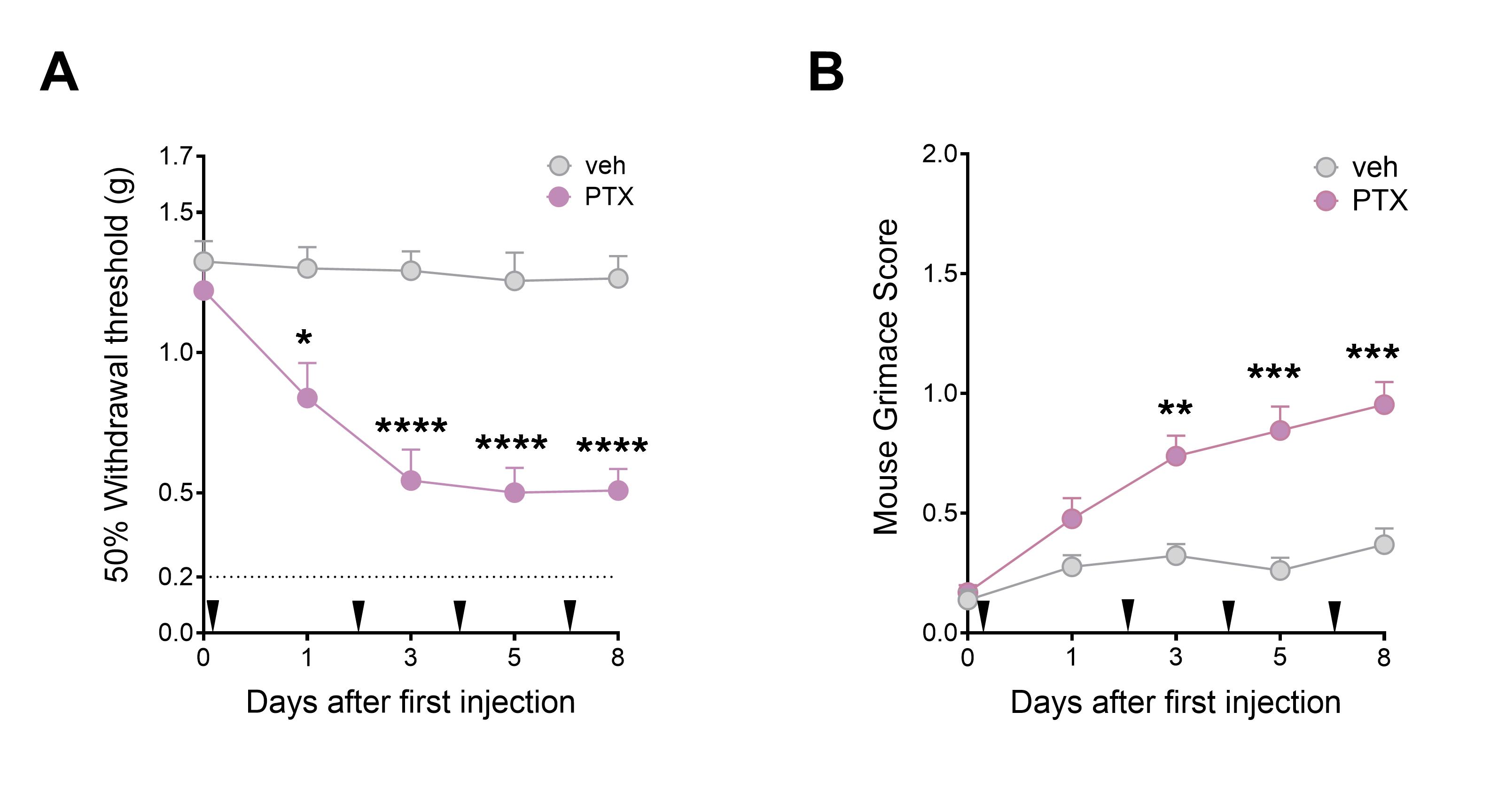

**Sup. Fig. 1.** Paclitaxel induces mechanical allodynia and spontaneous pain in mice. **A**. Time course of mechanical sensitivity in WT mice at 1, 3, 5 and 8 days after PTX (i.p.) injection. **B**. Time course of grimacing score in WT mice at 1, 3, 5 and 8 days after PTX (i.p.) injection. Data are presented as the mean ± SEM. *p < 0.05, *p < 0.01, ****p < 0.0001 (n = 13 animals per group, with 7 male and 6 female mice) as determined by two-way RM ANOVA followed by Bonferroni’s test. Arrow heads show the time of PTX injection
